# Degenerative changes are associated with severity of anterior cruciate ligament injury within the skeletally immature joint

**DOI:** 10.1101/2022.11.12.516262

**Authors:** Danielle Howe, Jacob D. Thompson, Stephanie D. Teeter, Margaret Easson, Olivia Barlow, Emily H. Griffith, Lauren V. Schnabel, Jeffrey T. Spang, Matthew B. Fisher

**Author notes:** **One Sentence Summary:** Joint degeneration was associated with degree of joint instability after partial and complete ACL injury in a skeletally immature porcine model.

## Abstract

Anterior cruciate ligament (ACL) injuries are a major problem in the pediatric and adolescent populations. Some of these injuries extend only partially through the tissue cross-section; yet, there is limited data to inform clinical treatment of such partial tears. In particular, it is unknown how injury severity impacts long-term degenerative changes in the joint. Here, we leverage a skeletally immature preclinical porcine model to evaluate joint biomechanics and degeneration after partial (isolated anteromedial (AM) or posterolateral (PL) bundle) or complete ACL injury. Six months after injury, joint laxity increases were minimal after PL bundle injury, minor after AM bundle injury, and major after ACL injury. Joint degeneration (evaluated in the cartilage and meniscus) was minimal after PL bundle injury, moderate after AM bundle injury, and substantial after ACL injury. With subjects grouped by clinical Lachman grade (indicating the extent of joint destabilization), degeneration was associated with increasing grade, irrespective of injury type. These findings point to the importance of considering joint laxity as a factor when treating young patients, particularly those with partial ACL injuries.

---

Injuries to the anterior cruciate ligament (ACL) of the knee, particularly those occurring in children and adolescents, are a significant clinical and societal concern (*1*). The total number of ACL injuries across all age groups is substantial (200,000-250,000 yearly in the United States alone) (*2, 3*). A specific concern is the 2-4 fold increase in the rate of injury and subsequent surgical treatment in the pediatric population compared to adults (*4–6*).

Preventing joint degeneration is a major concern. The natural occurrence of joint degeneration after ACL rupture is well documented in adults (*7*). In the pediatric population, full-thickness ACL tears result in two-fold increases in knee laxity (*8*) and 3-4 times greater risk of developing meniscal and cartilage lesions with delayed surgical intervention (*9*). It is commonly assumed that joint instability after ACL injury is a major factor in subsequent joint degeneration (*10–12*), though the degree of joint instability required to cause degeneration is unknown for skeletally immature patients. Nevertheless, surgical reconstruction is the standard practice to restore joint stability for young, active patients with complete ACL tears (*13*), despite concerns about growth plate disruption.

For partial-thickness ACL tears, which account for 10-27% of diagnosed ACL injuries (*14*) and result in a more minor (∼30% on average) increase in knee laxity (*15*), the superiority of operative or non-operative treatment to limit degenerative changes is less evidence-based. On one hand, subsequent meniscal and chondral pathologies after non-surgical treatment of partial tears are reported in young patients (*16, 17*). On the other hand, the invasive nature of reconstructive surgery can exacerbate degenerative changes (*18*), and approximately 50% of young patients who undergo ACL reconstruction still develop osteoarthritis after 10 years (*19*).

A major complicating factor in determining the best clinical treatment is that not all partial tears are the same. The ACL has two major subregions, the anteromedial (AM) bundle and the posterolateral (PL) bundle, and partial tears can occur in either. Tear location may impact functional outcomes after partial injury, and the results may differ between children and adults. Specifically, a study in adult patients reported similar joint stability after isolated AM and PL bundle injuries (*15*), while a study in pediatric patients reported better outcomes for AM bundle injuries (*20*). Furthermore, in a skeletally immature pig model, the biomechanical function of the AM and PL bundles of the ACL depends on age and sex, such that the AM bundle plays a larger functional role in older versus younger subjects and males versus females (*21, 22*). Since evidence regarding treatment of bundle-specific partial injuries in children and adolescents is sparse, there is a need for further investigation of long-term outcomes and degenerative changes in skeletally immature subjects. Large exploratory studies in humans are challenging to perform in the pediatric patient population. However, the pig is a valuable preclinical model with similar ACL anatomy and function to humans (*23–25*), which can be used to control injury severity and assess functional outcomes and degenerative changes after injury.

In the present study, we investigated biomechanics, remodeling, and degeneration in the knee joint after isolated AM bundle, PL bundle, and complete ACL tears in a juvenile porcine model. We hypothesized firstly that partial (AM and PL bundle) tears would result in small levels of instability and degeneration relative to complete ACL tears. Secondly, we hypothesized that remaining soft tissues in the joint would remodel in response to changes in biomechanical loading after ACL injury, such that the degree of remodeling would associate with injury severity. At 6 months after injury, joint laxity showed no change after PL bundle injury, minor increases after AM bundle injury and major increases after ACL injury. Similarly, PL bundle injury resulted in minimal degeneration, AM bundle injury resulted in moderate degeneration, and complete ACL injury led to substantial degeneration, such that the degree of degenerative changes was associated with joint instability.

## Results

### Joint kinematics after partial and complete ACL injury

To evaluate the impact of AM and PL bundle tears on knee function and degeneration in skeletally immature subjects, we performed arthroscopic transection and removal of the AM bundle (AM-), PL bundle (PL-) or ACL (ACL-) in juvenile Yorkshire crossbreed pigs and assessed biomechanics and joint health after 24 weeks (Fig. 1a). Joint kinematics were evaluated using a robotic testing system first under applied anterior-posterior (AP) drawer to simulate clinical exams (Fig. 1b-e). AP tibial translation (APTT) was greater in AM-joints (46%, P<0.0001) and ACL-joints (195%, P<0.0001) compared to contralateral controls, while PL-joints showed no significant difference (P=0.22) (Fig. 1c-d). Between injury groups, the inter-limb difference in APTT was significantly different for all comparisons (P≤0.0061) (Fig. 1e). Under applied compression (Fig. 1f-i) (to simulate weight bearing) and varus-valgus moments (Fig. 1j-m) (to simulate rotational loads), anterior tibial translation (ATT) was only greater in ACL-joints compared to contralateral controls (327%, P=0.0005) (Fig. 1h), and the interlimb difference in ATT was greater for the ACL-group compared to AM- and PL-groups (P<0.0001 for each) (Fig. 1i). Likewise, varus-valgus (VV) rotation was greater in ACL-joints compared to contralateral controls (33%, P=0.012) (Fig. 1l), with a larger interlimb difference compared to AM- and PL-groups (P≤0.0037) (Fig. 1m). Sex did not significantly affect kinematics under AP or VV loads, but under compression, ATT was significantly greater in males than females within the AM-group (P=0.003) (Supplementary Fig. 1). Overall, PL bundle injury induced minimal joint instability, AM bundle injury induced moderate joint instability, and complete ACL injury induced major joint instability.

**Fig. 1.**
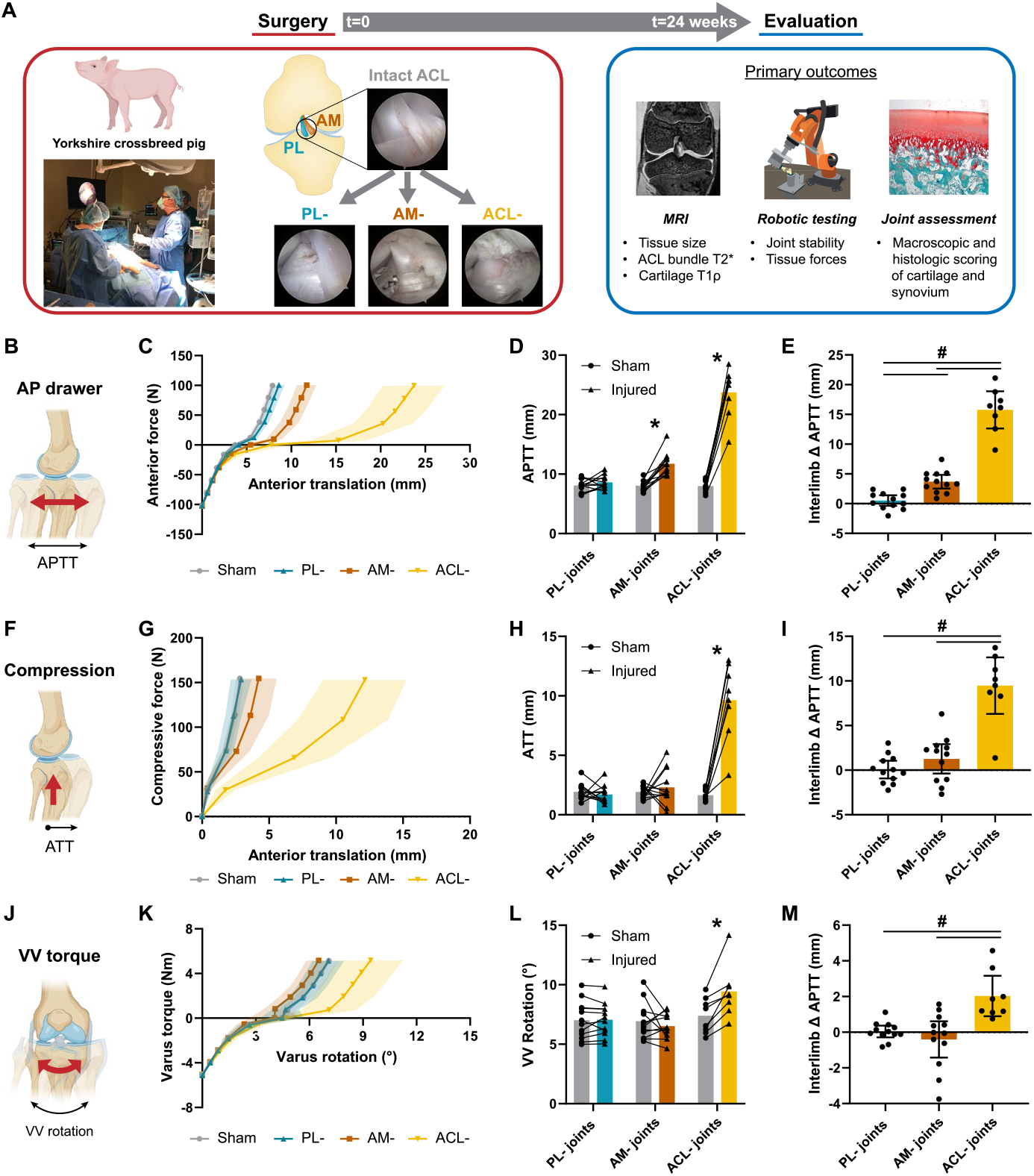
Biomechanical assessment of porcine joints 6 months after partial and complete ACL transection. a, Schematic of arthroscopic transection of the AM bundle, PL bundle, or entire ACL, followed by evaluation after 24 weeks. b, Diagram of anterior-posterior (AP) loads applied to the tibia. c, Resulting mean force-translation curves. AP tibial translation (APTT) shown as d, absolute values and e, interlimb differences. f, Diagram of compressive load applied to the tibia. g, Resulting mean force-translation curves. Anterior tibial translation (ATT) shown as h, absolute values and i, interlimb differences. j, Diagram of varus-valgus (VV) torques applied to the tibia. k, Resulting mean torque-rotation curves. VV rotation shown as l, absolute values and m, interlimb differences. Red arrows indicate the direction of the applied load, and black arrows indicate the direction of measured translation (b, f, j). Points and shaded area represent mean ± 95% CI (c, g, k). Lines connect paired joints (d, h, l). Individual data points (n=12 for PL- and AM-groups, n=8 for ACL-group) presented with mean ± 95% CI (e, i, m). * P<0.05 between injured and contralateral, from paired t-test with Holm-Sidak correction. #P<0.05 between injury type, from one-way ANOVA with Tukey post hoc test.

### Subhead 2: Remodeling of the uninjured bundle of the ACL

In partial injury (AM- and PL-) joints, the in-situ force (Fig. 2a) carried by the remaining ACL bundle was greater compared to the corresponding bundle force in contralateral controls under applied anterior drawer (P≤0.00019 for both injury types) (Fig. 2b). However, under joint compression, the remaining bundle force was elevated only for AM-joints (P=0.0016) (Fig. 2c). Remaining bundle in-situ force did not differ on average under varus or valgus moments, although considerable specimen-to-specimen variability existed (Supplementary Fig. 2). To assess remodeling of the ACL that may drive functional changes, the cross-sectional area (CSA) of the remaining bundle in injured joints was measured from MRI and compared to the corresponding bundle in contralateral joints (Fig. 2d). The remaining bundle was larger in CSA compared to contralateral controls for PL-joints (AM bundle) by 13% and AM-joints (PL bundle) by 60% (P≤0.022, for both) (Fig. 2e). In addition, bundle collagen organization was assessed through T2* mapping using MRI (Fig. 2f). No differences were found in median T2* value of the remaining bundle in injured joints compared to contralateral controls for either injury group (Fig. 2g).

**Fig. 2.**
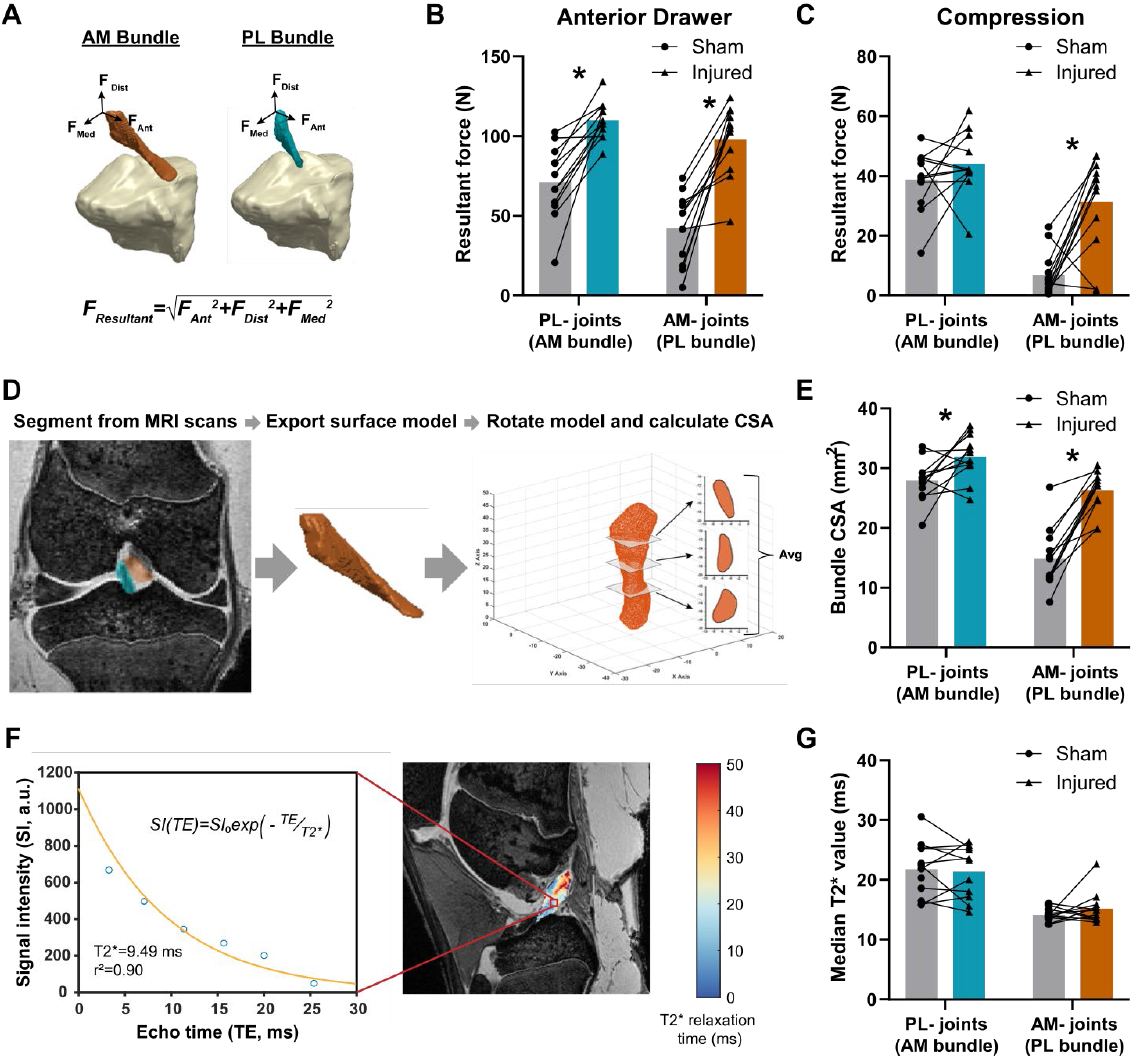
Remodeling of the remaining bundle after transection of the AM or PL bundle of the ACL. a, Schematic of in-situ bundle forces measured under applied loads. Resultant force in the uninjured bundle under applied b, anterior drawer and c, compression. d, Schematic demonstrating methods used to measure bundle cross-sectional area (CSA) from segmented MRI scans. e, CSA of the uninjured bundle. f, Schematic showing representative sagittal MR image overlaid with the T2* map of the ACL. Plot shows signal intensity at the six measured echo times (blue circles) and fitted monoexponential decay curve (yellow) for a representative voxel. Calculated T2* relaxation time and goodness-of-fit (r^2^) shown in graph. g, Median T2* value of the uninjured bundle. Individual data points (n=12 per injury group) presented with mean/median. Lines connect paired joints. *P<0.05 between injured and contralateral, from paired t-test with Holm-Sidak correction.

### Subhead 3: Cartilage and synovium degeneration

To assess if the type of injury impacted degeneration of articular cartilage in the joint, T1ρ mapping was performed via MRI, and the median T1ρ value was calculated for the medial and lateral femoral and tibial articular cartilage (Fig. 3a). Higher T1ρ values are linked to lower proteoglycan content (*26, 27*). T1ρ values relative to the contralateral control were not significantly elevated in any compartment for the PL-group, but were elevated in the medial femoral and lateral tibial compartments for the AM-group and the lateral femoral and lateral tibial compartments for the ACL-group (P≤0.022 for all, Fig. 3b-e). Furthermore, the interlimb difference in T1ρ values were significantly greater in the AM-group relative to the PL-group for the medial femoral compartment and in the ACL-group relative to the PL-group for the lateral femoral and lateral tibial compartments (P≤0.012 for all).

**Fig. 3.**
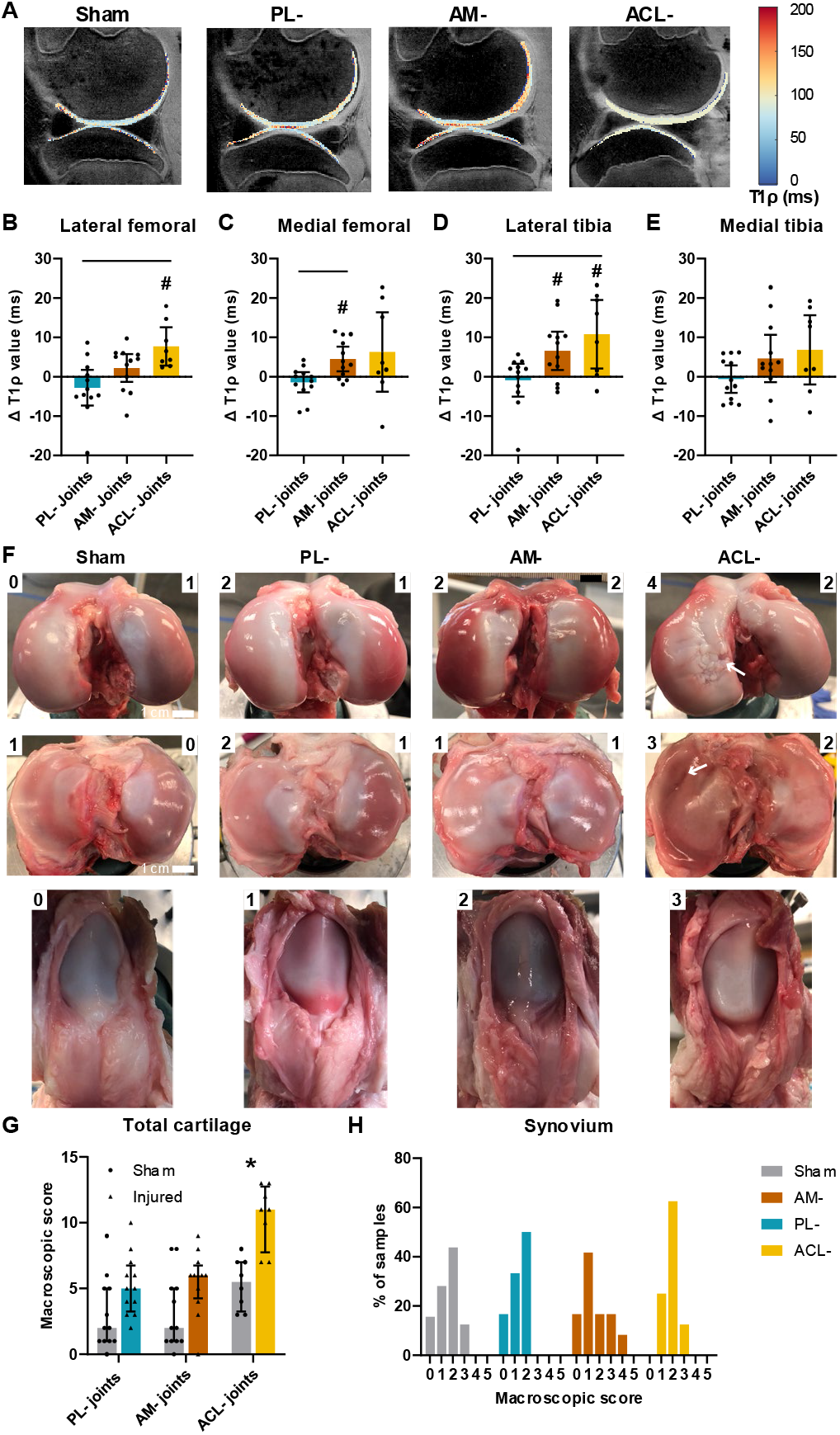
Cartilage degeneration and synovium inflammation. a, Representative sagittal MR images overlayed with T1ρ maps of articular cartilage. Interlimb differences in T1ρ value for b, lateral femoral, c, medial femoral, d, lateral tibial, and e, medial tibial articular cartilage. #P<0.05 compared to 0 from one sample t-test; bars indicate P<0.05 between injury type, from one-way ANOVA with Tukey post hoc test or Welch ANOVA with Dunnett post hoc test for data sets with unequal variance. f, Representative images of articular cartilage and synovium, with macroscopic scores shown in the corner of each image. White arrows indicate cartilage erosions reaching the subchondral bone. g, Composite macroscopic OARSI scores summed from the four articular cartilage compartments. *P<0.05 compared to contralateral controls from Wilcoxon matched-pairs test. h, Histogram of macroscopic OARSI scores for synovium, presented as a percent of samples for each group. Individual data points (n=12 for PL- and AM-groups, n=8 for ACL-group) presented with mean ± 95% CI (b-e) or median ± IQR (g).

Macroscopic articular cartilage damage was scored according to Osteoarthritis Research Society International (OARSI) guidelines (*28*) for the four compartments (Fig. 3f, Supplementary Fig. 3) and summed for a composite joint score. The composite joint scores were significantly elevated only the ACL-group compared to contralateral controls (P=0.016 Fig. 3g). Synovium pathology was similarly evaluated through macroscopic scoring, though no significant differences were found between groups (Fig. 3h). A small subset of cartilage was evaluated histologically, but there was high sample-to-sample variability and little discernable differences between injury groups. (Supplementary Fig. 4). Overall, minimal cartilage degradation was observed in PL-joints, and similar increases in cartilage T1ρ values (an early indicator of cartilage degradation) were observed in AM- and ACL-joints; however, macroscopic cartilage degeneration was elevated only in ACL-joints.

### Subhead 4: Meniscus degeneration

The biomechanical effect of injury on the menisci of the knee joint was assessed by measuring the in-situ load carried by each tissue under the applied loading conditions. The load carried in the medial meniscus under applied anterior drawer was 33% greater for AM-joints and 3.7-fold greater for ACL-joints compared to contralateral joints (P≤0.029 for each) (Fig. 4a), while no significant differences were measured under applied compression (Fig. 4b). The load carried in the lateral meniscus was greater compared to contralateral joints for AM-joints under applied anterior drawer (by 2.7-fold, P=0.008, Fig. 4c) and for ACL-joints under applied compression (by 2.1-fold, P=0.045, Fig. 4d). No differences in meniscal loads were measured in PL-joints relative to contralateral controls. Next, the volume of the menisci were measured from MRI (Fig. 4d) and compared between injured and control joints. The medial and lateral menisci were 11% and 7% larger in AM-joints and 28% and 30% larger in ACL-joints, respectively (P≤0.048 for all, Fig. 4f-g). No changes in meniscal size were measured in PL-joints. For ACL-joints, in addition to the measurable increases in size, the menisci qualitatively developed an oblique shape, and medial menisci developed an unorganized fibrous posterior outgrowth, evident macroscopically and on MRI (Supplementary Fig. 5). Overall, both in-situ loading and volume of the menisci showed no change in PL-joints, moderate increases in AM-joints, and substantial increases in ACL-joints.

**Fig. 4.**
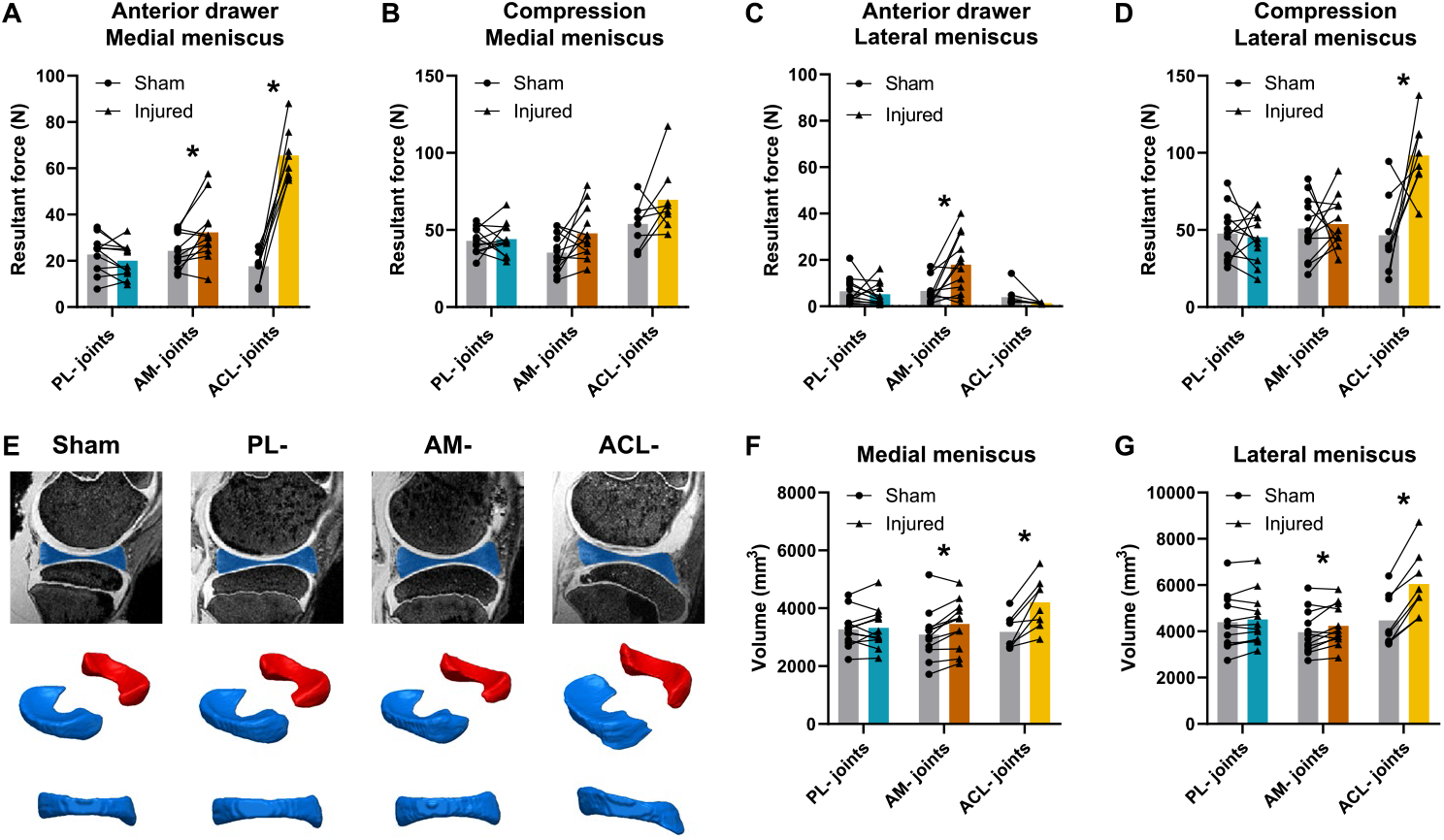
Changes in meniscal loading and size after partial and complete ACL transection. In-situ forces in the medial meniscus under applied a, anterior drawer and b, axial compression. In-situ forces in the lateral meniscus under applied c, anterior drawer and d, axial compression. e, Representative 3D models of menisci from each injury group, generated from MRI scans. Medial menisci shown in blue, lateral menisci in red. Volume of the f, medial meniscus and g, lateral meniscus. Individual data points (n=12 for PL- and AM-groups, n=8 for ACL-group) presented with mean. Lines connect paired joints. *P<0.05 between injured and contralateral, from paired t-test with Holm-Sidak correction.

### Subhead 5: Clinical implications and associations

To investigate whether findings were similar for partial injuries at different ages during skeletal growth, AM and PL bundle injuries were created in adolescent (6-month-old) pigs, and similar results were found (See Supplemental Results and Supplemental Figures 6-7). Using pooled data from both age groups, changes in joint laxity following injury were associated with degenerative changes in the cartilage. Linear regressions were run between interlimb differences in APTT and degenerative outcomes (Fig. 5a-h, Supplementary Fig. 8). Across injury types, significant positive associations existed between interlimb differences in APTT and mean side-to-side differences in T1ρ of the lateral femoral cartilage (r^2^=0.16, P=0.0089), lateral meniscus volume (r^2^=0.73, P<0.0001), and medial meniscus volume (r^2^=0.47, P<0.0001) (Fig. 5a,e, Supplementary Fig. 8). Within injury type, however, there were generally weaker associations for lateral femoral cartilage T1ρ and the interlimb APTT difference (r^2^=0.001-0.24) (Fig. 5b-d). For medial meniscus volume, poor associations were also found between the changes in medial meniscus volume and interlimb APTT increases within PL-injured (P=0.58) and ACL-injured (P=0.42) joints. The strongest positive association was found between the medial meniscus volume change and APTT in AM-injured joints, though again with an r^2^ value of only 0.26 (P=0.046, Fig. 5f-h, Supplementary Fig. 8m-p). Similarly, no significant associations were observed for changes in medial femoral cartilage T1ρ values or lateral meniscus volume and changes in APTT for any injury group (Supplementary Fig. 8e-l).

**Fig. 5.**
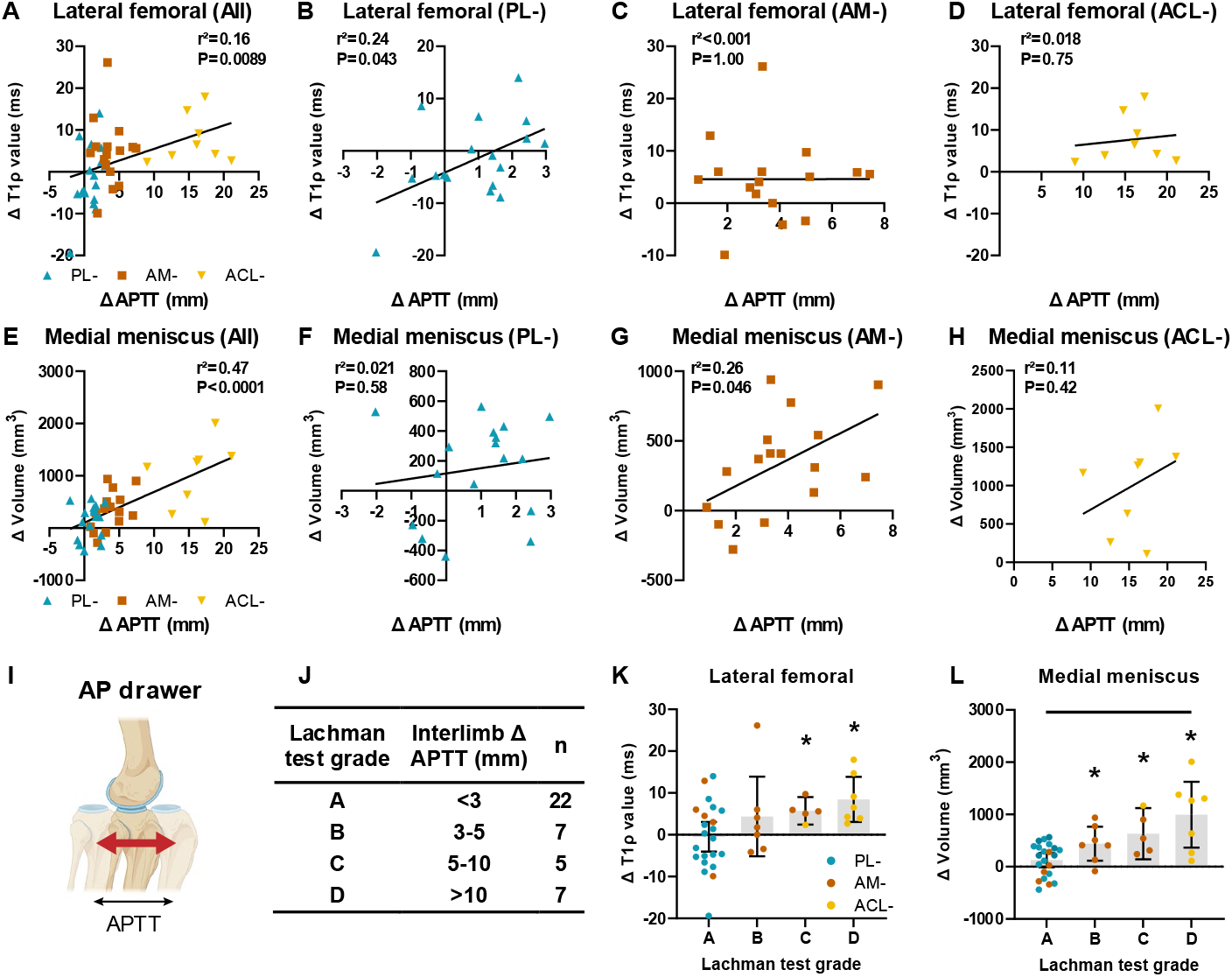
Degenerative outcomes as a function of changes in laxity. a-h, Linear regressions of interlimb difference in APTT versus (a-d) lateral femoral joint cartilage T1ρ and (e-h) medial meniscus volume, across injury groups and separated by PL-, AM-, and ACL-injured joints for pooled juvenile and adolescent pigs. i,j, Lachman test grade from applied anterior-posterior drawer in biomechanical testing used to separate interlimb difference in k, lateral femoral joint cartilage T1ρ and l, medial meniscus volume. *P<0.05 compared to 0 from one sample t-test; bars indicate P<0.05 between injury type, from one-way ANOVA with Tukey post hoc test. Statistical results from linear regression analyses shown in graph. Individual data points (n=17 for PL- and AM-groups, n=8 for ACL-group) presented with mean ± 95% CI (k-l).

To place the observed degenerative outcomes in a clinically relevant framework, specimens were grouped by equivalent Lachman test grade (Fig. 5i-l, Supplementary Fig. 9), which is a common clinical exam to assess joint stability after ACL injury. Generally, mean side-side differences in lateral femoral cartilage T1ρ values (P=0.048) and medial (P=0.0002) meniscus volume increased progressively for increasing Lachman test grades. Within grade, significant interlimb differences were measured for lateral femoral cartilage T1ρ values only for specimens with Lachman test grades C and D (P≤0.0086), but significant increases in medial meniscus volume were measured for specimens with grade B, C, and D (P≤0.023 for each). Linear regressions were performed within Lachman grades to further investigate associations between joint stability and degeneration (Supplementary Fig. 9), but these were generally weak (r^2^<=0.18). Pooling all injury types, these findings indicate that joint destabilization is positively associated with the extent of joint degradation, but strong associations were not readily apparent within an individual injury type or clinical test grade level.

## Discussion

There is presently a lack of data to inform clinical treatment of partial ACL injury in young patients, and large exploratory clinical studies are difficult to perform in this patient population. The present study provides a comprehensive analysis of functional and degenerative changes in the knee joint after partial (isolated AM or PL bundle) and complete ACL injury in skeletally immature subjects using a preclinical porcine model and clinically relevant outcome measures. Our results indicate that ACL injury severity (partial or complete) impacts outcomes, and equally importantly, that partial injury location (AM or PL bundle) also impacts outcomes. Across all injury types, the degree of degenerative changes was strongly associated with the extent of joint destabilization. Joint laxity increases were minimal for PL bundle injury, minor for AM bundle injury, and major for ACL injury. In both partial injury cases, the remaining bundle remodeled through an increase in CSA to attempt to restore ACL function and stabilize the knee joint. Cartilage degeneration and meniscal hypertrophy were observed to a minor degree after AM bundle injury and to a major degree after ACL injury, but not after PL bundle injury. Generally, sex did not impact degenerative outcomes in this model.

The joint laxity findings in this study are consistent with studies in humans, which report minor (0-6 mm) increases in laxity after partial ACL tears (*8, 15, 18, 20*) compared to major (6-12 mm) increases in laxity after complete ACL tears (*8, 15*). ACL injuries are commonly diagnosed on the field or in the office through physical examination using a Lachman test, which assesses tibial translation under anterior loads, or a pivot shift test, which evaluates tibial subluxation under compressive and rotational loads (*29*). Therefore, the small differences in laxity for partial ACL injuries point to challenges in diagnosis, particularly for PL bundle tears. This is supported by a clinical study in adults which found that 50% of partial tears confirmed through arthroscopy were diagnosed as a false negative using an arthrometer designed to quantify joint laxity (*30*). In the current study, no significant difference in side-to-side laxity was measured under anterior or rotational loads for the PL bundle injury group. In fact, all specimens from the PL bundle injury group fell within the Lachman grade A (indicative of a “normal” joint), suggesting that many of these injuries may go undiagnosed and untreated. Imaging, primarily MRI, can add value as a complementary diagnostic tool in cases where physical examination is inconclusive. Therefore, physical exams paired with MRI appear to be the best current process for diagnosing partial ACL tears, although there remains space for improvement of diagnostic sensitivity and specificity (*15, 31*). Diagnostic accuracy has the potential for improvement through implementing higher resolution MRI protocols or newer sequences capable of better contrast for ligaments (*32*). There also remains the potential for development of new clinical exams which can detect partial injuries.

Our functional data is consistent with clinical studies in adults which reported that a greater number of PL bundle injuries were biomechanically functional than AM bundle injuries (*15, 33–35*). Data is less definitive in pediatric patients, since currently only two studies evaluate outcomes after partial injury in young patients (*18, 20*). Kocher et al. found patients with a PL bundle injury were more likely to require a subsequent surgery (*20*), which seems contradictory to findings in this study. However, survivor bias may be a source for this discrepancy. In that study, patients with Lachman grade C or D underwent surgical reconstruction and were not included, which would exclude more AM bundle injuries than PL bundle injury based on our laxity findings. Additionally, it is possible that a greater percentage of AM bundle tears progress to complete ACL tears before receiving treatment. Even without progression to a complete tear, our data shows that degenerative changes can occur after partial ACL injury with joint instability, suggesting that surgical intervention may be of value even in cases of minor joint instability (Lachman Grade B or higher). Conversely, the lack of degenerative changes in partial ACL injury with no joint instability could provide a rationale for non-operative treatment, allowing more growth to occur prior to surgical intervention if needed.

Our findings are consistent with preclinical in-vivo studies of partial ACL injury as well. A study in skeletally mature sheep found gross and histological cartilage degradation and meniscal damage 40 weeks after AM bundle injury (*36*). Another study in skeletally mature rabbits found histological evidence of meniscal degeneration 8 weeks after both AM and PL bundle injuries (*37*). In adult rabbits, partial (“medial half”) and complete ACL injuries led to an increase in joint laxity and histological cartilage damage, and histological scoring was associated with anterior joint stiffness (*38*). Although factors including skeletal maturity, injury location, and time from injury differ between these studies and our study, the consistent findings show that partial ACL injury alone can lead to long-term joint degeneration across different preclinical small and large animal models. Furthermore, our study extends these prior findings to show that degeneration occurs even in the context of skeletally immature joints, despite the potential for higher healing/remodeling capacity.

Several animal studies have tried to associate joint instability and degeneration (*38–40*). However, these studies often involve creating injuries to multiple ligaments (*39*) or stabilizing the joint via surgery (*40*), which limits clinical relevance. Moreover, no study spans the full spectrum of instability corresponding to Lachman grades A through D. This is crucial since patients with grades A or B would be most likely to be treated non-surgically in humans.

Additionally, degenerative outcomes are often assessed via histology, which limits their clinical translatability. In our work, we comprehensively compare different types of ACL injury that create a full range of joint instability from Lachman grades A through D, allowing a comprehensive association to degeneration. Additionally, we leverage clinically relevant outcome measures (MRI) that could be used to motivate confirmatory studies in humans (though we also performed traditional gross and histological scoring).

Along these lines, the cartilage degeneration that we measured macroscopically and from MRI did not necessarily correspond to one another, particularly in the AM bundle injured group which showed significantly elevated cartilage T1ρ values but little macroscopic cartilage damage. Furthermore, histological scoring of the cartilage was highly variable and did not show clear associations with macroscopic scoring or MRI T1ρ relaxation values. These metrics inherently measure different outcomes, with macroscopic scoring reflecting large structural damage and loss of tissue, T1ρ values reflecting the matrix composition of existing cartilage tissue, and histological scoring reflecting local, 2D structural and cellular damage. While elevated T1ρ values are often used as an early predictor of future degeneration (*41*), macroscopic and histological scoring reflects later-stage structural degeneration. These metrics may capture different phases of degeneration, and we would predict that macroscopic scores for the AM bundle injury group would increase at a longer follow-up timepoint. The difference between outcome metrics also highlights the limited 2D nature of histology (only looking at a thin slice) and suggests that 3D metrics provide a clearer picture of overall cartilage degeneration.

In addition to cartilage, changes in the meniscus were also associated with joint instability, suggesting meniscal hypertrophy may also be an early indicator of joint degeneration. This is supported by data from a guineapig model, where meniscal hypertrophy was tied to cartilage damage after ACL injury (*42*). Although little to no data exist regarding meniscal size changes after ACL injury in humans, larger meniscus volume in middle-aged women was tied to later development of structural OA (*43*). While it is common to evaluate meniscal tears after ACL injury, meniscal size, independent of the presence of injury, is often overlooked. However, meniscus size may be an important indicator of overall joint health. Furthermore, meniscus size may be a more repeatable metric across time and would not require specialized MRI sequences, such as T1ρ.

The current work has implications for treatment of partial ACL tears, since we showed significant meniscal and chondral pathology after partial ACL injury at the equivalence of a grade B or higher Lachman test. Although we saw differences in outcomes due to injury type, the extent of destabilization may be a better metric as a basis for treatment, since we saw significant associations between cartilage degeneration and joint laxity even within AM bundle injuries. Our results suggest that surgical intervention may be appropriate for patients with Lachman grade B or above, and more likely to be necessary in patients with an AM bundle tear rather than PL bundle tear. Additionally, the increase in remaining bundle CSA with no change in T2* value indicates that viable tissue capable of functional remodeling can remain after a partial ACL injury. This points to advantages in surgical techniques which preserve the intact tissue. Single bundle augmentation techniques which repair or reconstruct only the injured bundle are relatively new but gaining in popularity (*44–48*). These techniques have the advantage of preserving the functional remaining tissue and may cause less donor site morbidity since smaller grafts would be required.

This study includes several limitations, including the use of an animal model. While the pig is an established preclinical model for musculoskeletal studies (*23*) and the ACL bundles function similarly between pigs and humans (*24*), there are some differences in knee anatomy (*25*) and physiological loading compared to humans. Furthermore, human patients are prescribed physical therapy and activity restrictions, while we did not attempt to restrict animal activity or acquire metrics for physical activity. Moreover, our evaluation of joint stability does not account for active stabilization provided by the muscles, so it is possible that the joint is more stable in-vivo. Nevertheless, the degenerative changes we detected occurred in the presence of muscle function in-vivo. Additionally, our surgical model of ACL injury does not account for damage to other tissues that may occur during the high impact of a naturally occurring ACL injury. However, our model does isolate the effects of progressive degeneration caused by the loss of joint stability. The injured bundle or entire ACL was removed in our surgical model. This prevents any healing, which has been observed in a limited capacity in clinical cases (*49*). An additional limitation includes the use of contralateral limbs as a control. These may differ from joints in healthy animals due to altered gait or limb compensation after injury of the ipsilateral joint. Lastly, the conclusions in this study are limited to only one timepoint after injury and do not capture the time course of degenerative changes. Future longitudinal studies to assess the progression of the degenerative outcomes would be useful for distinguishing early predictors of degeneration and identifying an ideal window for implementing treatment interventions to prevent degenerative changes.

This work fills key gaps in knowledge about degenerative changes after partial and complete ACL injury in skeletally immature subjects. We leverage a large animal model to collect detailed data about joint biomechanics, tissue remodeling, and cartilage degeneration which are not possible to collect in clinical studies. Our data show that the extent of destabilization after an ACL injury may serve as a predictor of degeneration, irrespective of the injury type (partial or complete) or partial injury location (AM or PL bundle). Associations in this study also imply that there may not be a key threshold beyond which degenerative changes occur, but instead, a continuous gradient where even small increases in laxity can cause minor degeneration. Overall, this study provides a fundamental basis for informing clinical treatments in young patients with partial ACL tears.

## Methods

### Subhead 1: Study design

The objective of this study was to evaluate the outcomes of partial ACL injury as a function of tear location (AM or PL bundle), with covariates of sex and age, in a skeletally immature surgical porcine model. All animals were bred at the North Carolina State University Swine Educational Unit, and all experimental protocols were approved by the North Carolina State University Institutional Animal Care and Use Committee. Sample sizes were calculated using a power analysis described in the “Statistical analysis” section. A total of 32 juvenile Yorkshire crossbreed pigs (age: 13 ± 1 week, weight: 34 ± 5 kg) underwent unilateral arthroscopic transection of the AM bundle (n=6 female, n=6 male), the PL bundle (n=6 female, n=6 male), or the complete ACL (n=4 female, n=4 male) with a sham operation performed on the contralateral joint. Injury type and injured limb side were randomized. To confirm findings in an older age group, a small subset of 9 adolescent pigs (age: 25 ± 1 week, weight: 129 ± 21 kg) underwent unilateral arthroscopic transection of the AM bundle (n=3 female, n=1 male) or PL bundle (n=4 female, n=1 male). One adolescent animal which was meant to undergo AM bundle transection was excluded due to a pre-existing partial ACL tear to the PL bundle. The endpoint was set at 24 weeks to experimentally evaluate joint biomechanics, cartilage health, and remodeling of soft tissues in the knee joint. Data were collected and analyses were performed in a blinded manner, where possible. No outliers were excluded in the presented data.

### Subhead 2: Surgical procedure

Prior to surgery, all pigs were sedated and received analgesics and antimicrobials. Operations were performed arthroscopically (Arthrex Synergy System) under general anesthesia. The knee joints were accessed through medial and lateral parapatellar incisions (∼0.5 cm long). The infrapatellar fat pad was partially removed using a 3.5 mm motorized shaver (SabreTooth, Arthrex) to visualize the ACL. A freer elevator was inserted between the AM and PL bundles near the tibial insertion and used to bluntly separate the bundles. A triangle knife (Ectra II Knife 4448, Smith & Nephew) was inserted into the separation and (1) turned anteriorly to transect the AM bundle, (2) turned posteriorly to transect the PL bundle, or (3) used to transect the full cross-section of the ACL. The remnant of the transected bundle or complete ligament was debrided using a shaver to ensure loss of function and prevent healing. In the contralateral sham-operated control limb, the infrapatellar pad was similarly partially removed. The ACL bundles were separated with a freer elevator but not transected. Following the operation, skin incisions were suture closed, a local analgesic was administered through subcutaneous injection at the portal sites, and the joint was covered with an antimicrobial adhesive drape. Animals were given analgesics and antimicrobials for 3 days post-operatively. Animals were allowed full weight bearing immediately following surgery and housed in pens with no restriction of movement. After 6 months (23.9 ± 0.3 weeks), all animals were euthanized, and hind limbs were collected. Skin and excess muscle were removed, and knee joints were wrapped in saline-soaked pads and stored at −20 °C.

### Subhead 3: Magnetic resonance imaging

Prior to scanning, joints were thawed. Joints were imaged at full extension. Imaging was performed using a Siemens Magnetom Prisma 3T MRI system and a knee coil. Joints were scanned using a T1ρ fast low-angle shot (FLASH) sequence (FOV: 128×128×84mm, voxel size: 0.4×0.4×1.5mm, FA: 10°, TR: 8.74ms, TE: 4.60ms, spin-lock frequency: 500 Hz, TSL: 0, 10, 20, 30, 40ms) and a T2 susceptibility weighted imaging (SWI) sequence (FOV: 160×160×128mm, voxel size: 0.4×0.4×0.8mm, FA: 12°, TR: 31.0ms, TE: 3.3,7.1,11.4,15.7,20.0,24.4ms). Following scanning, joints were stored at −20 °C for subsequent biomechanical testing. T1ρ and T2* relaxation maps were created by fitting monoexponential decay function on a voxel-by-voxel basis to the 5 spin-lock times and 6 echo times, respectively, using a custom MATLAB script.

Articular cartilage from the medial and lateral condyles of the femur and tibia were manually segmented by a single user from T1ρ images with 0 ms spin-lock time using commercial software (Simpleware ScanIP). Using a MATLAB script, the segmented areas were transposed onto T1ρ relaxation maps described above, and the median T1ρ relaxation time was calculated for each of the four segmented cartilage compartments.

Similarly, the AM bundle, PL bundle, complete ACL, and medial and lateral menisci were manually segmented by a single user from T2* images with 15.7ms echo time. The median T2* relaxation time was calculated for each ACL bundle. Additionally, the volume of each segmented tissue was recorded, and the cross-sectional area (CSA) of the AM and PL bundles and the complete ACL were calculated as previously described (*21, 22*). Briefly, each segmented tissue was exported as a surface mesh, and analyzed using a MATLAB script. Each mesh was rotated such that a line of best fit through the mesh vertices aligned with the z-axis. The CSA was calculated for slices of the mesh in 1 mm increments along the z-axis, then averaged over the middle 50% of slices.

### Subhead 4: Biomechanical testing

Biomechanics of the knee (stifle) joint were evaluated using a 6 degree-of-freedom (DOF) robotic testing system (KR300 R2500, KRC4, Kuka) equipped with a universal force-moment sensor (Omega160 IP65, ATI Industrial Automation), and integrated through the simVitro software knee module (Cleveland Clinic). To prepare for testing, joints were thawed at room temperature and trimmed to expose the femoral and tibial diaphyses. The femoral head and the distal end of the tibia were removed using a bone saw, then bones were fixed using fiberglass-reinforced epoxy (Everglass, Evercoat) in molds designed to fit in custom clamps for testing. Joints were again stored at −20 °C and thawed at room temperature one day prior to robot testing.

The femoral end of the joint was fixed to a rigid testing platform, while the tibial end was fixed to the robotic system at 40° of flexion. A 3D digitizer (G2X MicroScribe) was used to determine an anatomic coordinate system. Joints were sprayed with saline and lightly wrapped in a pad throughout testing to maintain hydration. A passive flexion path was determined by flexing the joint from 40° to 90° while minimizing forces/moments in the remaining 5 DOFs. Then, loads/torques were separately applied to the tibia in the anterior-posterior (AP) (100 N), proximal (axial compression, 140 N), and VV (5 Nm) directions/rotations at 60° of joint flexion while recording kinematic paths. Kinematics were then replayed while recording forces. Next, the joint capsule was removed, and kinematics were replayed while recording forces. The difference in forces measured between the intact and capsule-deficient condition was calculated as the force carried by the joint capsule, by the principle of superposition (*50*). This process was repeated after transection of the AM bundle (or any remnant), PL bundle (or any remnant), MCL, LCL, PCL, medial meniscus, lateral meniscus, medial femoral condyle, and lateral femoral condyle. Joint laxity was measured by anterior-posterior tibial translation (APTT) under AP drawer, anterior tibial translation (ATT) under axial compression, and VV rotation under VV moments. Load-transformation curves were plotted for each loading condition using data points collected at intervals of 20% of the maximum applied load.

### Subhead 5: Macroscopic scoring

During robotic testing and tissue dissection, gross images were taken of the synovium following joint capsule dissection, as well as the femoral and tibial articular cartilage, following removal of all other tissues, during robotic testing. Synovium and cartilage were macroscopically scored using Osteoarthritis Research Society International (OARSI) guidelines for sheep and goat (*28*). Although are no scoring guidelines for pigs, this scoring system has been used to evaluate cartilage in previous porcine studies (*51, 52*). Synovial pathology was scored from 0 (normal) to 5 (severe), and articular cartilage of medial and lateral condyles of the femur and tibia were scored from 0 (normal) to 4 (large erosions down to subchondral bone) by two blinded reviewers. Disagreements between users were re-examined to determine a consensus. Scores from the four cartilage compartments were added to give a total joint score.

### Subhead 6: Histological analyses

Immediately following robotic testing, 5 mm coronal slabs were cut from the lateral femoral condyle (1.5-2 cm from the femoral notch apex) and lateral tibial plateau (1.5-2 cm anterior to the posterior boundary of the tibial plateau) using a bone saw. Samples were fixed in 4% paraformaldehyde solution at 4°C for 72 hours, then decalcified in Formical-2000 for 17 days. Samples were trimmed to 2-3mm in thickness and embedded in paraffin. 8 μm sections were taken from each sample and stained with safranin-O/fast-green. Whole sections were imaged at 4x magnification using an Olympus IX73 microscope with CellSens software. Two blinded reviewers scored samples adapted from OARSI guidelines (*53*) in the following categories: matrix staining, fibrillation, and focal cell loss.

### Subhead 7: Statistical analyses

Sample sizes were determined using a power analysis based on prior work that demonstrated the ability of transection of the AM bundle and ACL in cadaveric joints to generate different levels of initial joint instability (*54*). Using an overall α level of 0.05 (accounting for multiple comparisons), a power of 0.8, and an effect size of 1.5 for the AM-group, a sample size of n=12 was selected to measure a difference between injured and sham-operated joints. For complete loss of ACL function, much larger effect sizes are expected (>3), so a sample size of n=8 was used for this group.

Normality of all data sets was confirmed using Shapiro-Wilk test, and equal variance between data sets was tested using Brown-Forsythe test. Joint kinematics, tissue forces under applied loads, tissue size metrics, and bundle T2* values were compared between injured and sham-operated joints using paired t-tests with Holm-Sidak’s correction for multiple comparisons. Outliers were removed from lateral meniscus forces under anterior drawer using the iterative Grubbs’ method with an alpha of 0.05. Interlimb differences in joint kinematics, cartilage T1ρ, and meniscal volumes were compared between injury types and ages of injury using a two-way ANOVA with Holm-Sidak post hoc comparison. Interlimb differences in macroscopic and histologic cartilage scores were compared between AM-injury and PL-injured joints in juvenile and adolescent pigs with Mann-Whitney tests. For data sets with unequal variance, Welch ANOVA with Dunnett post hoc comparison was used. To evaluate clinically relevant outcomes, samples were grouped into equivalent Lachman test grades according to IKDC guidelines (A: <3 mm, B: 3-5 mm, C: 5-10 mm, D: >10 mm) (*55*), based on joint translations under AP drawer. Interlimb differences in cartilage T1ρ, macroscopic cartilage scores, and meniscal volume were compared between Lachman grades using one-way ANOVA and compared to zero with one sample t-test. Associations were assessed between interlimb differences in APTT and interlimb differences in cartilage T1ρ and meniscal volume using linear regression across all injury types and within injury types. All statistical analyses were performed in Prism (GraphPad), with overall significance set at α=0.05 for all analyses with corrections for multiple comparisons done as needed. Adjusted P values are reported. Data presented as mean ± 95% CI, unless stated otherwise, and data are reported in supplemental material.

## Data availability

The main data supporting the results in this study are available within the paper and its Supplementary Information.

## Acknowledgments

We would like to thank Laboratory Animal Resources (NC State) and the Biomedical Research Imaging Center (UNC-CH) for their contributions to this work. This work was supported by the National Institutes of Health (NIH) grants F31AR077997 and R01AR071985, and the National Science Foundation (NSF) grant DGE-1746939.

## Author contributions

D.H., L.V.S., J.T.S. and M.B.F. conceptualized the work. D.H., J.D.T., S.D.T., M.E., E.H.G., L.V.S., J.T.S. and M.B.F. designed the methodology. D.H., J.D.T., S.D.T., M.E., O.B., L.V.S., J.T.S. and M.B.F. investigated the work. Formal analysis was carried out by D.H., J.D.T., S.D.T., M.E., E.H.G. and M.B.F. Data was curated by D.H., J.D.T., S.D.T., and M.E. D.H., J.D.T., M.E. and M.B.F. created the figures for this work. Funding was acquired by D.H., J.D.T., L.V.S., J.T.S. and M.B.F., and the project administration was D.H. and M.B.F. with M.B.F. providing supervision. D.H. wrote the original draft and all authors reviewed and edited the paper.

---

**Supplementary Fig. 1.**
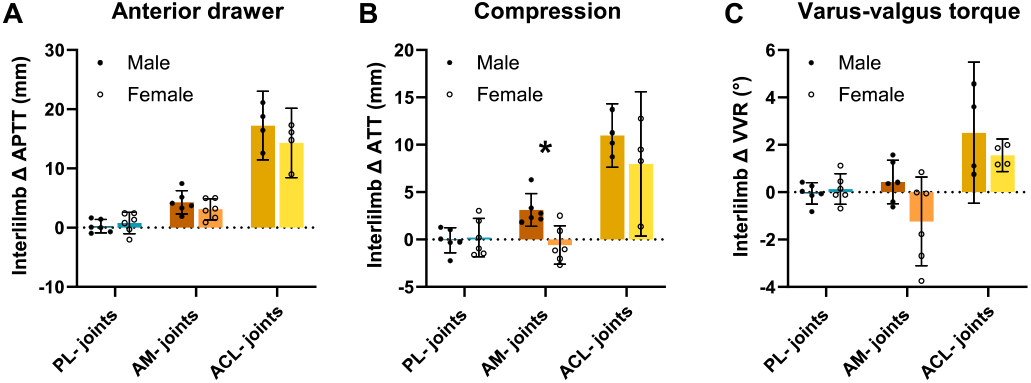
Sex-specific biomechanical outcomes. Interlimb differences in (A) anterior-posterior tibial translation (APTT) under applied AP drawer, (B) anterior tibial translation (ATT) under applied axial compression, and (C) varus-valgus (VV) rotation under applied VV moments. Individual data points presented with mean ± 95% CI. * P<0.05 between male and female, from one-way ANOVA with Tukey post hoc test.

**Supplementary Fig. 2.**
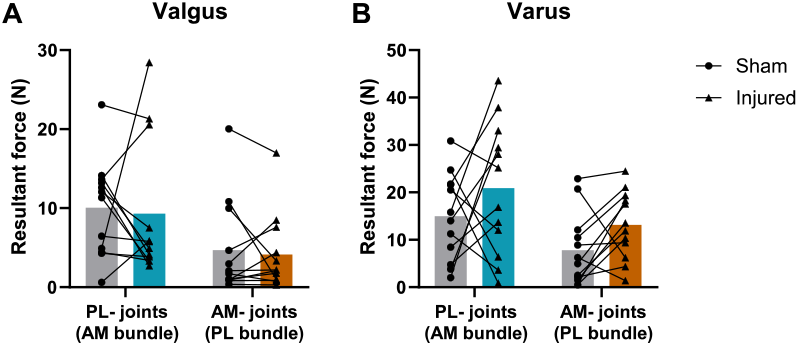
In-situ bundle forces under rotational loads. Resultant force in the uninjured bundle under applied (A) valgus moments and (B) varus moments. Lines connect paired joints. Individual data points presented with mean.

**Supplementary Fig. 3.**
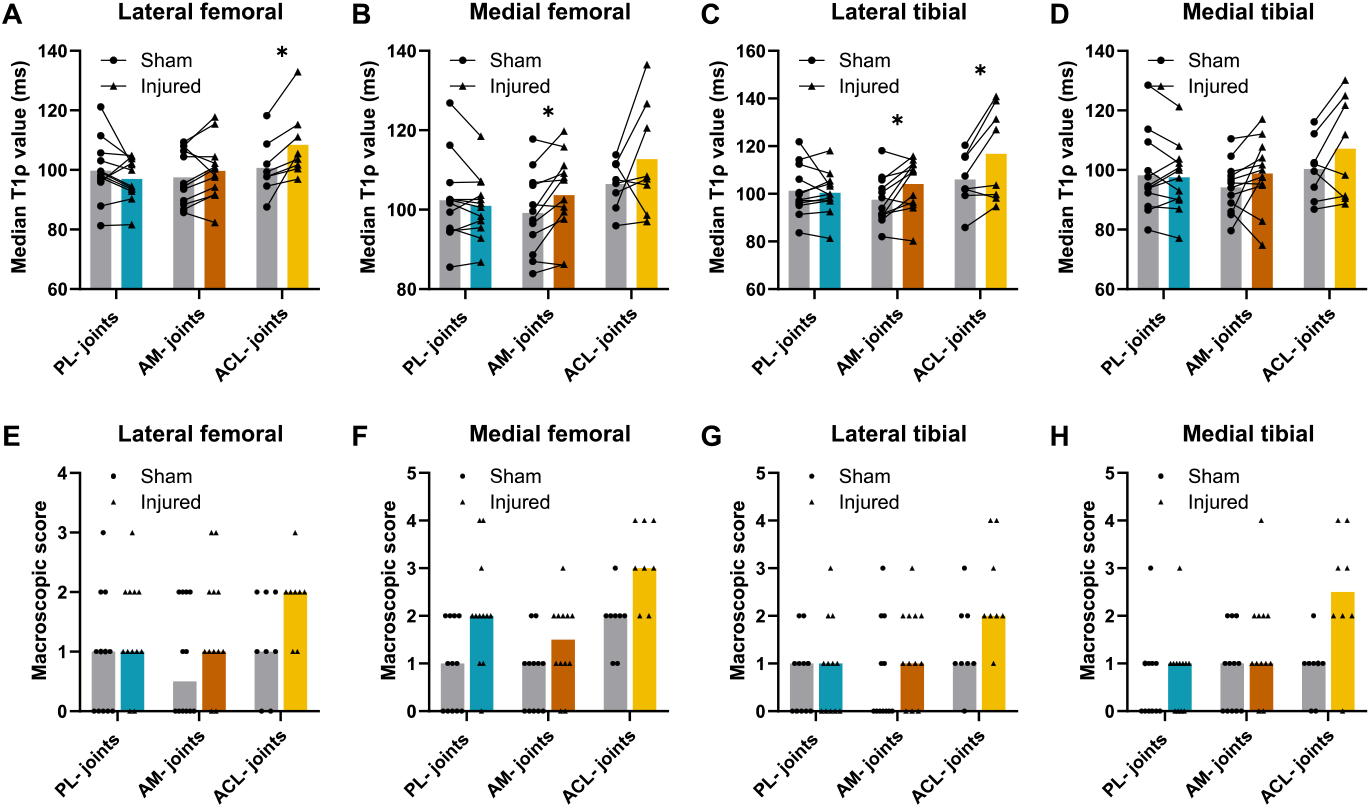
T1ρ values and macroscopic scores of cartilage compartments. T1ρ values of the (A) lateral femoral, (B) lateral tibial, (C) medial femoral, and (D) medial tibial cartilage compartments. Macroscopic OARSI score of the (E) lateral femoral, (F) lateral tibial, (G) medial femoral, and (H) medial tibial cartilage compartments. Individual data points presented with mean with lines connecting paired joints (A-D) or median ± IQR (E-H). *P<0.05 between injured and contralateral, from paired t-test with Holm-Sidak correction.

**Supplementary Fig. 4.**
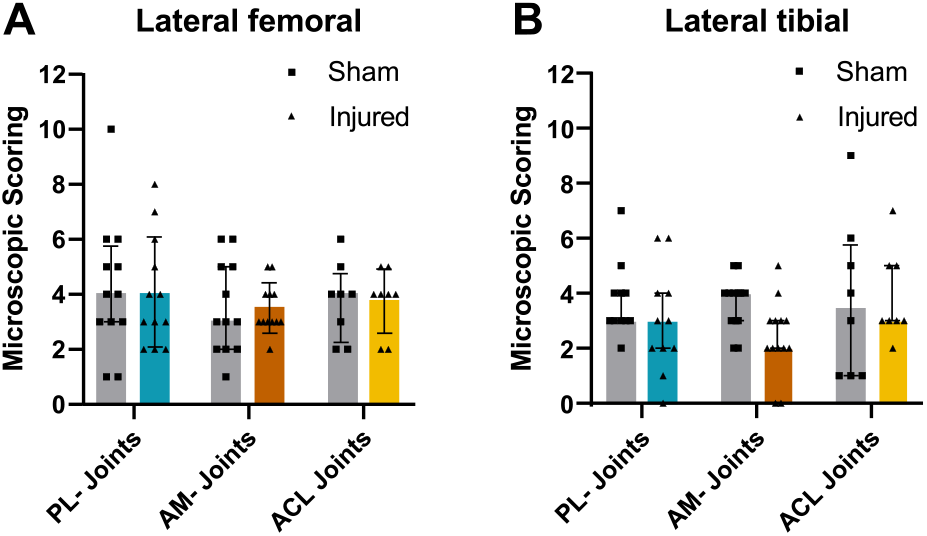
Histological scoring of lateral femoral and tibial cartilage samples stained with safranin-O and fast green. Representative images taken from samples in sham-operated, PL-, AM-, and ACL-joints. Summed composite histological scores shown for the (A) lateral femoral cartilage and (B) lateral tibial cartilage. Individual data points presented with median ± IQR.

**Supplementary Fig. 5.**
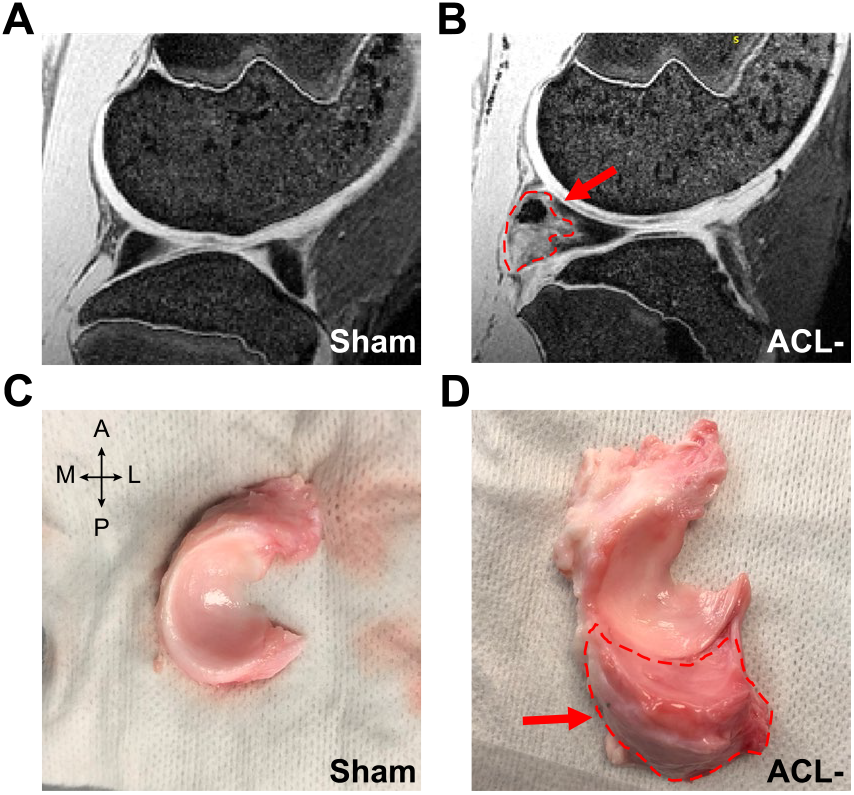
MRI and macroscopic images of medial menisci from sham-operated and ACL-joints. (A,C) Sham-operated control shows normal anatomy of the medial meniscus, where (B,D) red dotted lines and arrows indicate degenerative, fibrous outgrowth in ACL-injured joints. Anterior-posterior and medial-lateral directions indicated.

## Supplementary Results & Discussion: Extension to adolescent animals

To investigate whether findings were similar for partial injuries at different ages during skeletal growth, AM and PL bundle injuries were created in adolescent (6-month-old) pigs. Joint biomechanics, cartilage health, and meniscus volume were evaluated after 24 weeks. Interlimb differences in joint laxity were similarly elevated for AM-joints under AP drawer (P<0.05) in both the adolescent and juvenile injury groups, with no significant difference in laxity for the PL-group (Supplementary Fig. 6A, Fig. S6A). Laxity results were also similar between the two age groups under varus-valgus torque and compression for the PL-injured joints (Fig. S6B-C and Fig. 1). Both juvenile and adolescent age groups had similar increases of ∼1-2 mm in laxity under compression for AM-injured joints, yet the adolescent group had a statistically significant increase in laxity where the juvenile group did not (Fig. S6B and Fig. 1). For both age groups, lateral and medial femoral cartilage T1ρ values and composite macroscopic cartilage scores appeared similarly elevated in AM-joints but not PL-joints, with the medial femoral cartilage T1ρ having statistically significantly higher (P<0.05) T1ρ values for AM-injured joints for both juvenile and adolescent age groups (Fig. 5B-D and Fig. S6F-J). Lastly, the interlimb differences in medial and lateral meniscus volume were elevated ∼5-10% after partial injury in adolescent animals similar to juvenile animals (Fig. S6D and E). Although sample size was limited for adolescent animals (n=4-5 per injury type), these data highlight similarities across two age groups, adding confidence to our findings in younger animals with larger sample sizes.

**Supplementary Fig. 6.**
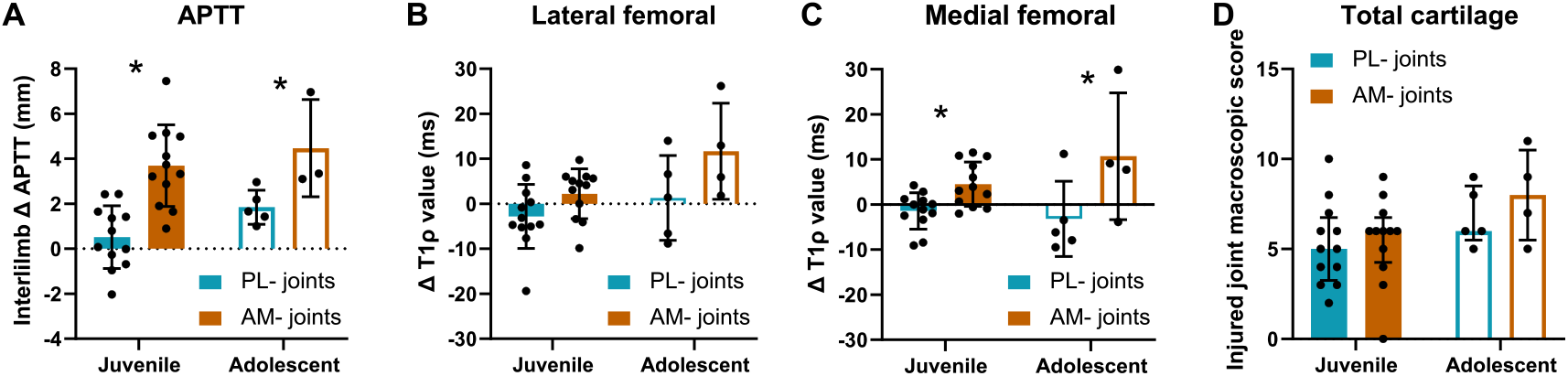
Joint laxity and tissue remodeling after partial ACL injury created in adolescent compared to juvenile pigs. (A) Interlimb differences in APTT under applied AP drawer. (B) Interlimb differences in T1ρ value for lateral femoral articular cartilage and (C) medial femoral articular cartilage. (D) Interlimb differences in macroscopic cartilage. Individual data points (n=5 for PL- and n=4 for AM-adolescent groups, n=12 for PL- and AM-juvenile groups) presented with mean ± 95% CI (A,B,D,E) or median ± IQR (C). *P<0.05 between PL- and AM-injured joints, from two-way ANOVA with Holm-Sidak correction. Multiple comparisons were run using Mann-Whitney test for interlimb differences in macroscopic scoring between PL- and AM-injured joints for juvenile and adolescent animals.

**Supplementary Fig. 7.**
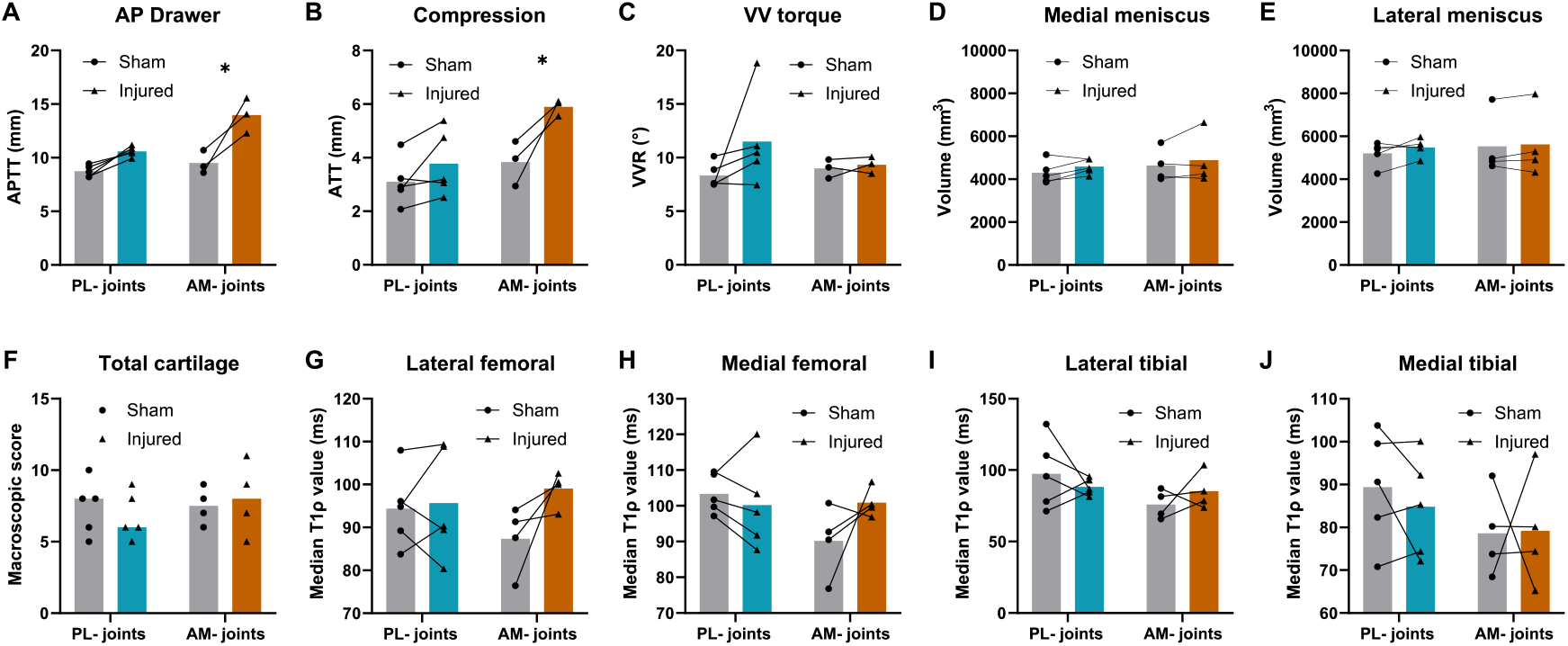
Biomechanical and degenerative outcomes after partial injuries created in adolescent (6 month) pigs. (A) APTT under applied AP drawer, (B) ATT under applied axial compression, and (C) VV rotation under applied VV moments. Volume of the (D) medial meniscus and (E) lateral meniscus. (F) Composite macroscopic OARSI scores summed from the four articular cartilage compartments. T1ρ value for (G) lateral femoral, (H) medial femoral, (I) lateral tibial, and (J) medial tibial articular cartilage. Lines connect paired joints. Individual data points (n=5 for PL- and n=4 for AM-groups) presented with mean. *P<0.05 between injured and contralateral, from paired t-test with Holm-Sidak correction.

**Supplementary Fig. 8.**
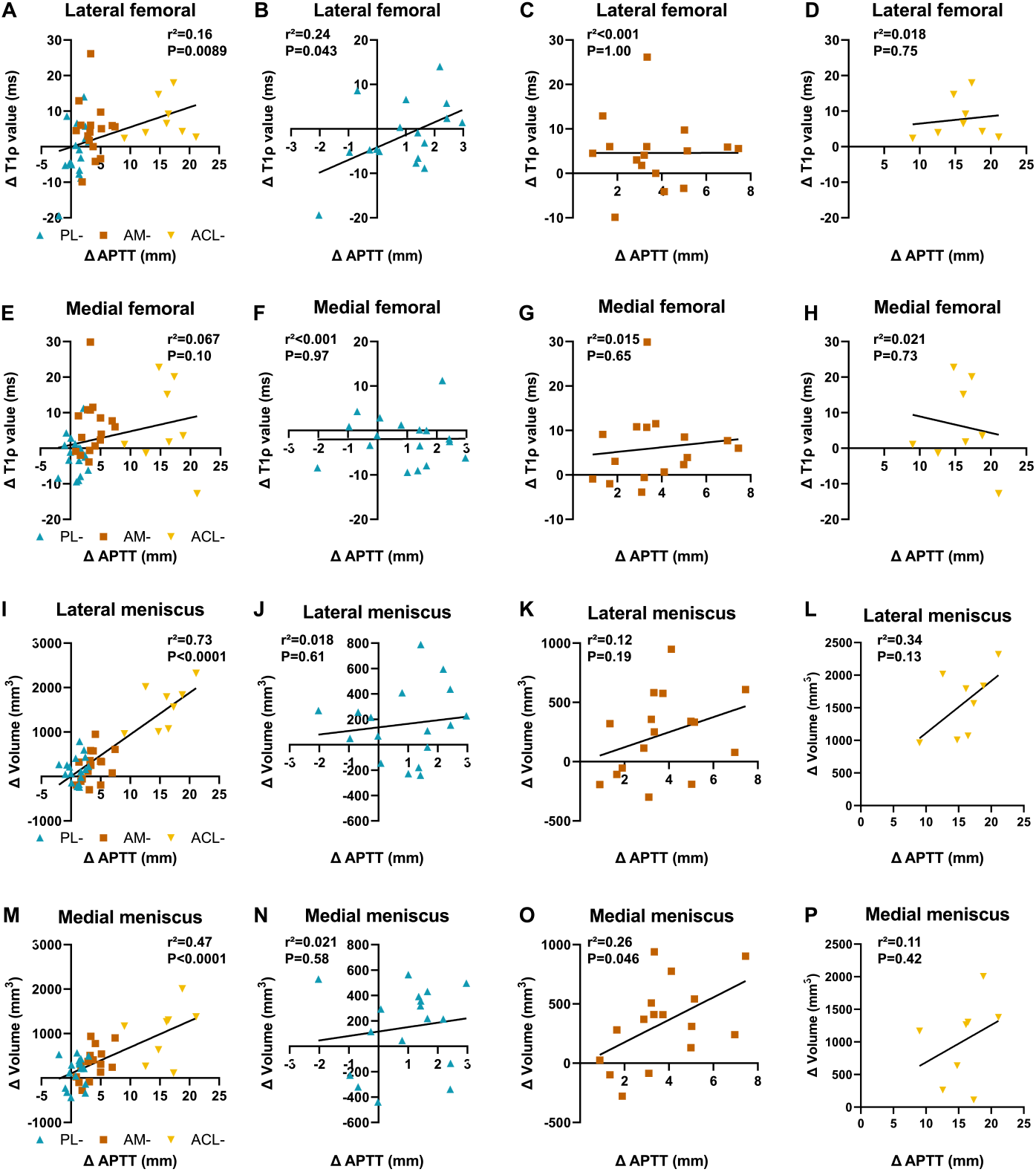
Interlimb difference in (A) lateral and (E) medial femoral cartilage T1rho values and (I) lateral and (M) medial meniscus volumes versus interlimb difference in APTT across all injury groups in pooled juvenile and adolescent pigs. Injury groups separated by PL-, AM-, and ACL-for interlimb differences in (B-D) lateral femoral cartilage T1rho values, (F-H) medial femoral cartilage T1rho values, (J-L) lateral meniscus volume, and (N-P) medial meniscus volume. Statistical results from linear regression analyses shown in graph.

**Supplementary Fig. 9.**
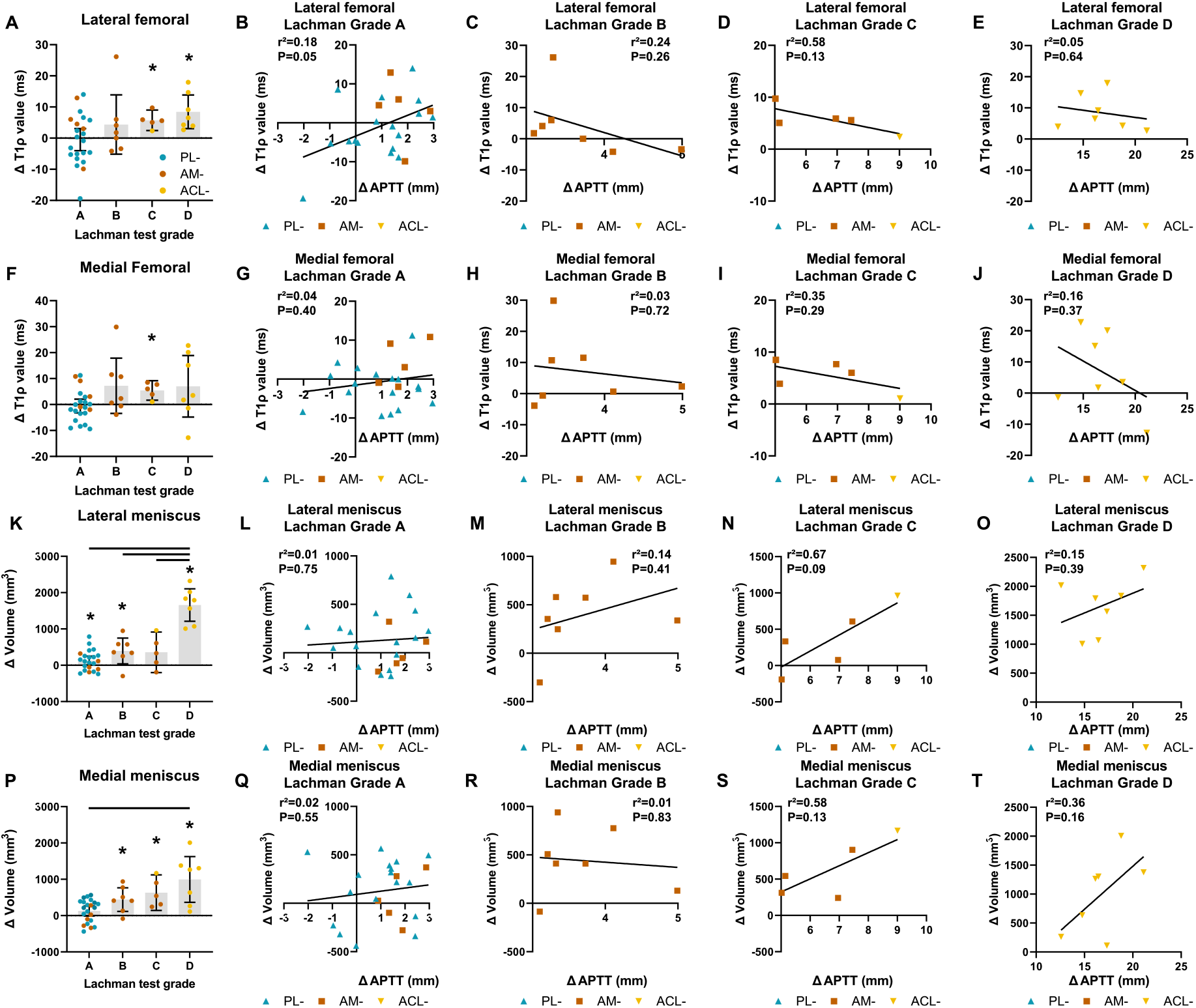
Interlimb difference in (A) lateral and (F) medial femoral cartilage T1rho values and (K) lateral and (P) medial meniscus volumes, grouped by the equivalent Lachman test grade in pooled juvenile and adolescent pigs. *P<0.05 compared to 0 from one sample t-test; bars indicate P<0.05 between injury type, from one-way ANOVA with Tukey post hoc test. Injury groups separated by Lachman grades A-D for interlimb differences in (B-E) lateral femoral cartilage T1rho values, (G-J) medial femoral cartilage T1rho values, (L-O) lateral meniscus volume, and (Q-T) medial meniscus volume. Statistical results from linear regression analyses shown in graph. Individual data points presented with mean ± 95% CI (A,F,K,P).

